# High-Resolution Epidemic Simulation Using Within-Host Infection and Contact Data

**DOI:** 10.1101/133421

**Authors:** Van Kinh Nguyen, Rafael Mikolajczyk, Esteban A. Hernandez-Vargas

## Abstract

**Background:** Transmission in epidemics of infectious diseases is characterized by a high level of subject-specific elements. These include heterogeneous infection conditions, time-dependent transmission potential, and age-dependent contact structure. These insights are often lost in epidemic models using population data. Here we submit an approach that can capture these details, paving the way for studying epidemics in a more mechanistic and realistic way.

**Methods:** Using experimental data, we formulated mathematical models of a pathogen infection dynamics from which we can simulate its transmission potential mechanistically. The models were then embedded in our implement of an age-specific contact network structure that allows to express all elements relevant to the transmission process. This approach is illustrated here with an example of Ebola virus (EBOV).

**Results:** The results showed that within-host infection dynamics can capture EBOV’s transmission parameters as good as approaches using population data. Population age-structure, contact distribution and patterns can also be captured with our network generating algorithm. This framework opens vast opportunities for the investigations of each element involved in the epidemic process. Here, estimating EBOV’s reproduction number revealed a heterogeneous pattern among age-groups, prompting questions on current estimates which are not adjusted for this factor. Assessments of mass vaccination strategies showed that a time window from five months before to one week after the start of an epidemic appeared to be effective. Noticeably, compared to a non-intervention scenario, a low vaccination coverage of 33% could reduce number of cases by ten to hundred times as well as lessen the case-fatality rate.

**Conclusions:** This is the first effort coupling directly within-host infection model into an age-structured epidemic network model, adding more realistic elements in simulating epidemic processes. Experimental data at the within-host infection are shown able to capture upfront key parameters of a pathogen; the applications of this approach will give us more time to prepare for potential epidemics. Population of interest in epidemic assessments could be modeled with an age-specific contact network without exhaustive amount of data. Further assessments and adaptations for different pathogens and scenarios are underway to explore multilevel aspects in infectious diseases epidemics.

## Background

Epidemics of infectious diseases are listed among the potential catastrophes and can be potentially misused as mass destruction weapons [1]. Overwhelming research efforts have been developed to early predict the danger of the epidemics but their crisis nature left scientists no better option than learning from the past [2, 1]. However, confronting outbreaks of emerging infections requires swift responses and thus the ability to evaluate quickly and early potential outcomes [1]. As such, computer simulations of epidemic models undoubtedly hold the potential as the first-aid toolbox for decision making amid the crisis [1, 3, 4].

A majority of epidemic modelling studies has exclusively relied on the availability of outbreak data [5, 6, 7, 8]. This approach requires that sufficient incidence data are available; for example, data at the end of an epidemic or at least until its peak [9]. As such, it would have limited applicability to newly emerging epidemics. Moreover, mechanistic models based on outbreak data are often oversimplified [10, 11]. For example, the effective transmission probability [7] has been usually simplified as a single parameter that reflects collective effects of the contact rate with the infectious, the infectivity of the infectious, and the susceptibility of the susceptible; influential factors of a disease transmission such as transmission probability and contact rate were mostly fixed while they dynamically change in reality [11]. As a result, these key processes in the disease transmission are lost, especially the transient nature of the infection course as well as the dynamics of the active population portion in an epidemic [12, 11].

In reality, the within-host infection process determines key parameters in the disease transmission [13, 12, 14, Fig. 1]. In an infected subject, interactions between the viruses and immune responses shape the viral load dynamics that ultimately defines the incubation period, the transmission potential, and the recovery rate [15, 14]. It is also evident that susceptibility to infection is not the same for all the susceptible but, among others, it is highly correlated with a subject’s age due to the aging of the immune systems [16, 17]. Differences in the within-host infection profile as well as the susceptibility to infection complicate greatly epidemic models but at the same time underline their influential roles in determining epidemics features and effects of certain intervention strategies [18, 19, 20].

**Figure 1.**
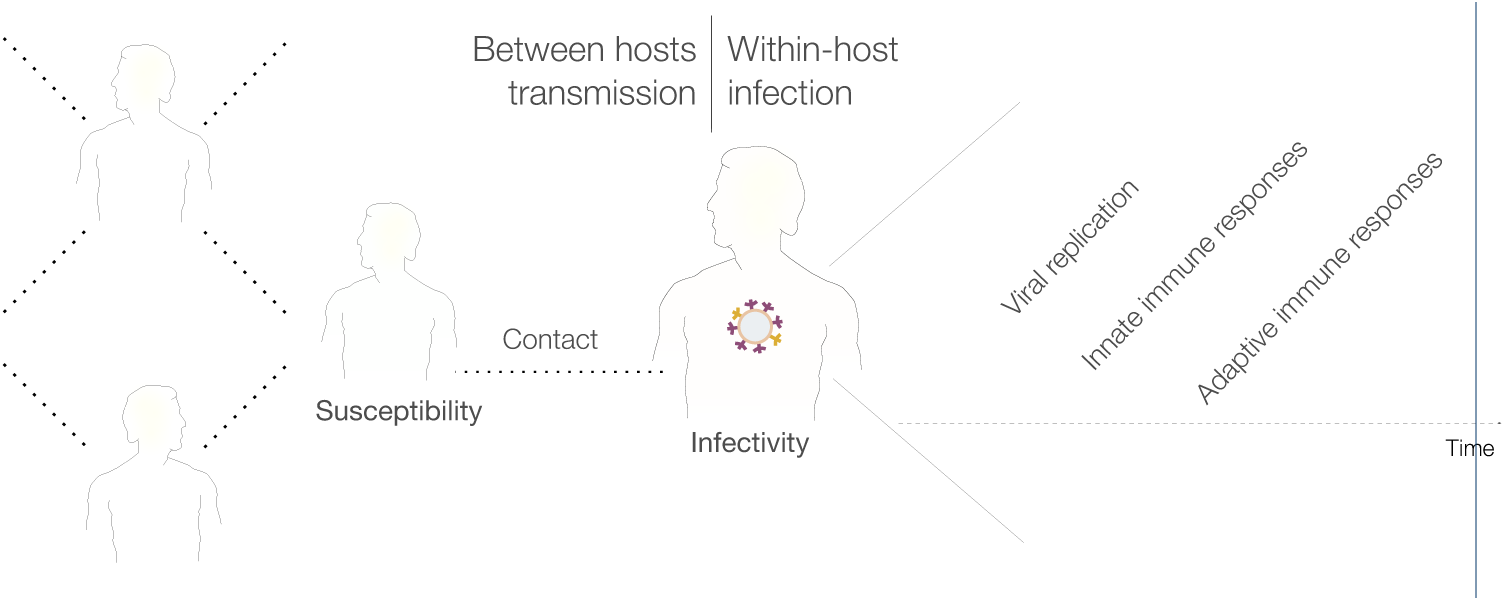
Schematic presentation of the two infections processes in an outbreak. At the within-host level, viral replication and immune responses race with each other that eventually determines an individual infectivity, for example, his symptoms and possibly behaviours. At the between hosts level, infected individuals make contact(s) with susceptible individual(s) that eventually lead to a transmission, depending on both the infectivity of the infectious and the susceptibility of the susceptible. Noting that while contact network can be fixed, the portion actively partake in epidemic spreading dynamically change over time [?, see]]Bansal:2007ct.

The interplays between within-host infection and between hosts transmission led to arising attempts connecting the two levels [14, 21, 22, 12, 23, 24, 25, 4], but the approach is still at its infancy [13]. On one hand, most of these models were conceptual and theoretical [13] or rely on assumptive and previously obtained parameter estimates [26, 15, 27]. We propose that this limitation can be overcome by using explicitly within-host infection model. On the other, implementations of population level models were either a general representation using probabilistic assumption [28, 29] or a demanding implementation of a particular population [27, 30, 15]. These approaches, while able to recover valuable insights, may not be representative and accessible for another population of interest, as none or massive amount data are needed. In this case, using measures of social mixing, such as number of contacts per day, can be highly consistent across regions [31, 32], representative for most connectivities relevant to disease spread [33, 34], and need not to be data-intensive [35].

Based on our previous studies of within-host EBOV infection [36, 37, 38], we brought forward a developed within-host infection model to study transmission fitness at the population level. We built a network model based on social contact data [31] and the respective epidemic simulation algorithm embedding the within-host model into the network model. In this way, the models are both data-informed while modest amount of data are needed. Parameters obtained from simulations were compared to those estimated based on actual outbreak data and empirical observations. The results showed that using with-host infection model not only uncovered faithful estimate of the transmission parameters, but also allowed the evaluations of detailed and realistic intervention effects. Implementations of the network model from social contact data is straightforward and can be extend for larger scale simulation on high performance computer clusters. In that capacity, epidemic assessments and preparations can be conducted quickly, ahead of time, and with high-resolution requirements.

## Methods

In an EBOV-infected subject, different immune systems components dynamically evolve in response to the viral replication dynamic. As a result, a series of events is triggered determining infection outcomes such as infectious status, symptoms, recovery, or death [39, 40, 41]. Therefore, the EBOV replication dynamics within a host were used in this paper to infer it transmission parameters.

### Within-host model

Using viral dynamics and immune responses data within a host, mathematical relations can be defined to test hypothesized infection mechanisms [42, 36]. In this context, non-human primates (NHPs) are the standard animal model for developing EBOV’s therapeutics and vaccines in humans [43, 44] which recently has been used to develop an effective vaccine against EBOV [45]. Epidemiological and pharmacological studies reported that a viral load higher than 10^6^ copies/mL [44, 46] is associated with a higher mortality rate, whereas observations on experimental data in NHPs showed that a viral load level higher than 10^6^ TCID_50_ was fatal [40, 39]. Here the viral load dynamics were simulated based on the model as follows [38]:

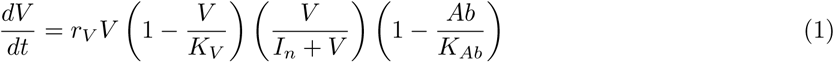

where *r*_*V*_, *K*_*V*_ and *I*_*n*_ denote the replication rate, the host’s carrying capacity, and a constraint threshold expressing the lag-phase growth of the virus. The parameter *K*_*Ab*_ represents the strength of the immune system at which the antibody titre inhibits the viral net growth rate [38]. The model parameters were obtained previously [38] using two experimental datasets on NHPs [39, 40]. The antibody dynamic (*Ab*) was also fitted in [38] to data of NHPs vaccinated with vesicular stomatitis virus (VSV-EBOV) vaccine [40]. The VSV-EBOV has recently showed efficacy in human [45]. Detailed of model fitting and data can be found in [38] with extracted parameters are also presented in Code and examples — Epidemic simulations.

### Simulated subject-specific infection course

To simulate subject-specific infection course, the antibody response strength *K*_*Ab*_ was varied from a normal level approximately 10^2.5^ [47, 40] to the highest observed level of 10^4.5^ [40]. This value was assumed to vary based on individual’s age assuming followed U-shaped function with higher susceptibility in infants and elderly [16, Fig. 1] (extracted data presented in Code and examples — Epidemic simulations). As infective dose can alter the course of infection [48], the initial condition *V*(0) of model Eq. (1) was varied depending on from whom a subject acquires the infection, i.e., equals the lethal dose (*V*_*c*_ = 10^0.15^ [38]) times the transmission potential of whom transmits the disease. Here we assumed a direct relation [13] between the transmission potential and the viral load at the time of infection, i.e., the transmission potential *p*_Trans_(*t*) = *V* (*t*)/*K*_*V*_. Note that *p*_Trans_(*t*) = 1 does not guarantee a successful transmission, but it was considered collectively with its contacts susceptibility and with the existence of such a contact (details in Code and examples — Epidemic simulations).

### Infection outcomes definitions

Empirical observations from both EBOV infected human and NHPs showed that the time from symptom onset to death is approximately one week [39, 40, 49]. Based on this and the viral load, we used the total area under the viral load curve (AUC) seven days post-infection in the subjects that died as a threshold above which the infection is lethal, i.e., 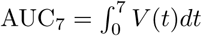. Otherwise, infected subjects were assumed recovered once the viral load was no longer detectable (Fig. 2). Depending on the infective dose and the adaptive immune response strength, an infection will manifest different viral dynamics. Based on that, we defined the transmission parameters as in Table 1A-C (detailed implementations can be seen in Code and examples — Epidemic simulations).

**Figure 2.**
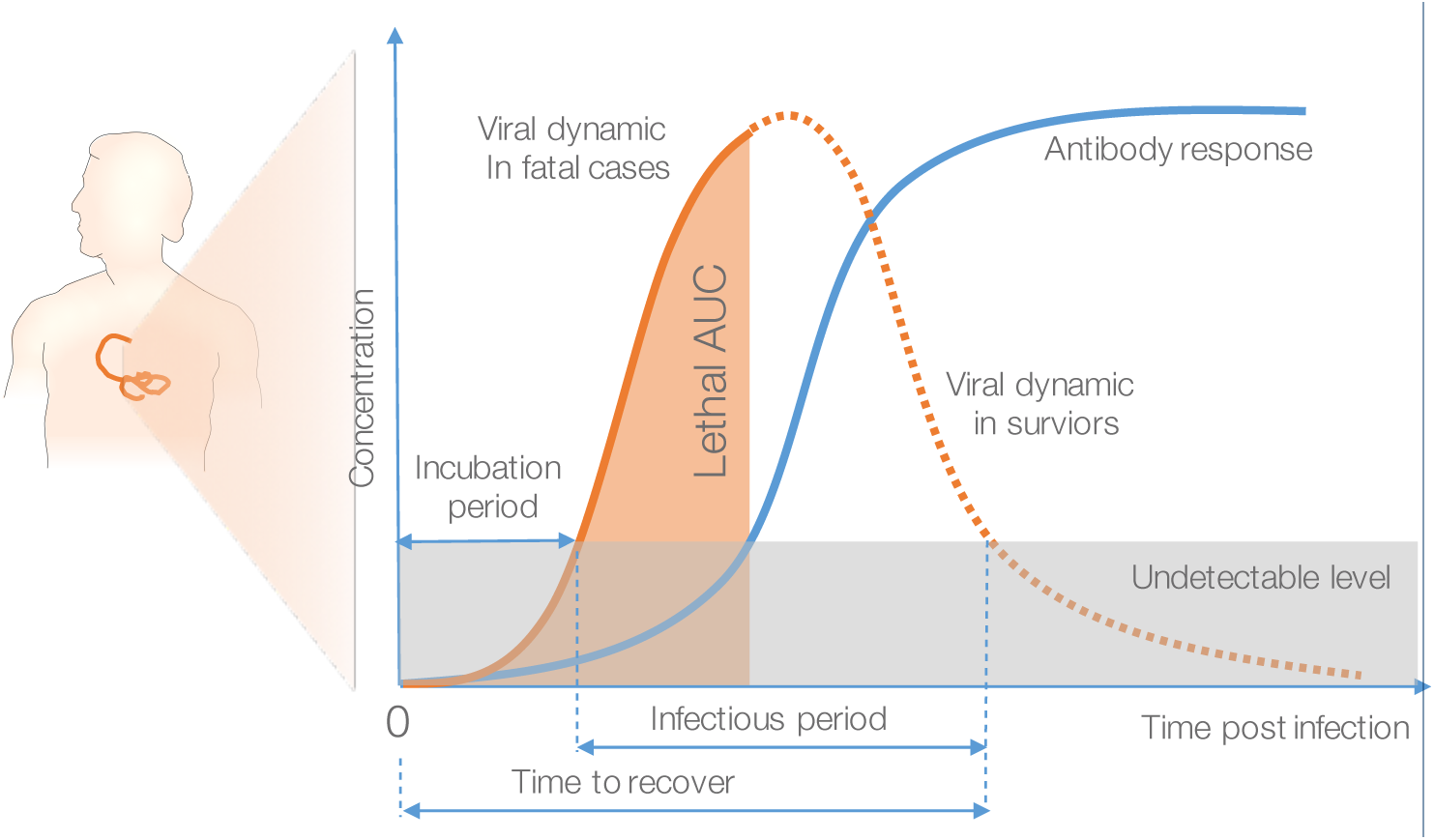
Simulated infection course using within-host infection dynamics. The viral replication, the antibody dynamics, and their interaction were modeled to define epidemiological parameters. It is assumed that when the EBOV-specific antibody concentration reaches certain threshold, it can inhibit the viral replication [50]. The total viral load under the curve (AUC) in lethal cases is used to define infection outcomes [51].

**Table 1.** Definitions of transmission parameters based on viral load and epidemics outcomes based on network model.

| Measure | Definition |
| --- | --- |
| A Incubation period | the interval between exposure to a pathogen and initial occurrence of symptoms [5] was defined from the infection time to the first time the viral load crosses over the detectable threshold (Fig. 2). |
| B Time from symptom onset to recovery[5] | defined as the interval between the first day of detectable viral load and the first day the viral load goes undetectable (Fig. 2). |
| C Time from symptom onset to death[5] | defined as the interval between the first day with detectable viral load and the day the area under the viral load curve (AUC) crosses the reference threshold $AUC_7$ (Fig. 2). |
| D Basic reproductive number ($R_0$) | calculated based on the network of infected subjects at the end of an epidemic. In terms of network models, this equals the mean degree distribution of the infected network, considering a directed network without loops (e.g., Fig. 5A). The $R_0$ by age-group was also calculated in the same fashion based on the assigned age-attribute. Note that in epidemics with intervention, the $R_0$ is called the *effective reproductive number* ($R_e$). |
| E Final infected fraction | the proportion of infected nodes at the end of the epidemic simulations. |
| F Case-fatality rate | the proportion of nodes died as a result of EBOV infection calculated as the end of epidemics. |

### The network model

The European’s contact patterns survey data [31] were used to generate a network model reflecting the number of contacts, the mixing patterns among age-groups, and a specific population age-structure. The age-distribution of the city Freetown in Sierra Leon was used as the reference [52]. A detailed description of the implementation can be found in Code and examples — Generating age-specific contact network. Because EBOV spreads through direct contacts with infectious subjects [48], and that the highest risk of infection is contacting with blood, faeces, and vomit [53], we used only the data of physical contacts and excluded those contacts with a duration less than five minutes. To account for the transmission route through funeral practices in EBOV outbreaks [2], we considered deceased EBOV-infected subjects infectious until they were buried. During the last epidemics in Sierra Leone, the time from death to burial was one to two days on average but can be a week [54]. As data regarding this variable’s distribution are not available, this number was randomly assigned using a truncated normal distribution at zero and seven with unit mean and variance (detailed implementations can be seen in Code and examples — Epidemic simulations).

### Transmission outcomes definitions

To obtain EBOV’s epidemics metrics, the within-host infection model was embedded into network model. Simulations of EBOV epidemic are detailed in Box 1. In short, a network of ten thousand nodes was generated. Scenarios in which the population was randomly vaccinated during one-week vaccination programs were tested and compared to a control simulation without interventions. For each scenario, one thousand simulations were performed, each of which started with a single random index case. Each time when a contact occurs, the viral load at the time point was extracted to determine the transmission potential. Next, the susceptibility of the contact persons were computed as a function of their age [16]. A Bernoulli trial was then used to determine if the contact results in an infection given the overall transmission probability. If the transmission succeeds, the newly infected subject has its own infection profile computed. Based on simulation outputs, the epidemic outcomes were determined as in Table 1D-F (detailed implementations can be seen in Code and examples — Epidemic simulations).

##### Box 1. EBOV epidemic simulation process

1. Choose index case(s), simulate infection course with lethal infective dose
2. For each time step *t*_*i*_ in simulation time *T*

a. For each infectious node i. Compute current transmission potential *p*_*I*_*i*__ = *V*( *t*_*i*_)/*K*_*V*_, *p*_*I*_*i*__ ∈ [0,1] ii. Find neighbours *j* = 0, …, *n*, compute their susceptibility with respect to their age [16]: *p*_*S*_*j*__ = *ƒ*_*S*_(age_*j*_), *p*_*S*_*j*__ ∈ [0,1] iii. Compute the transmission probability *p*_*i,j*_ = *p*_*I*_*i*__*p*_*S*_*j*__, decide if a neighbour is infected with a Bernoulli trial iv. Update newly infected nodes: store infection course, infection time, dose of infection, and fate decision. v. Update previously infected nodes if they are recovered or died.
b. Update death nodes: if buried removed the node from infectious nodes. Stop the simulation when there is no longer an infectious node. Noting that extra weights, e.g., contact duration, can be incorporated into iii.

### Computational implementation

The simulations were written in vectorized R language [55]. Computation of infection dynamics of newly infected node were done on-the-fly after obtaining its infective dose and immunization status. For nodes with identical conditions, their infection courses are copied instead of recomputing the ordinary differential equations (ODEs) for speed (Code and examples — Epidemic simulations). Repeated runs of epidemic simulations to obtain uncertainty estimates were done on computer clusters of the Center for Scientific Computing (CSC) of the Goethe University Frankfurt. Distributing of computation resources was sent from within R to SLURM Workload Manager.

## Results

### Basic transmission characteristics

Simulations of the outcomes of the within-host infection model showed a highly skewed distribution of the basic transmission parameters (Fig. 3). The incubation period derived from viral load dynamics ranged from 2.6 to 12.4 days (median: 3.8) compared to the previous estimates based on actual outbreak data ranging from 3.35 to 12.7 days [5]. The delay time from infection to recovery ranged from 6.9 to 17.6 days (median: 9.7) while previous estimates of this interval ranged from 2 to 26 days (median: 10) [5]. The time from infection to death ranged from 8.1 to 15.1 days (median: 9) compared to previous estimates ranged from 3 to 21 (median: 9–10) [5].

**Figure 3.**
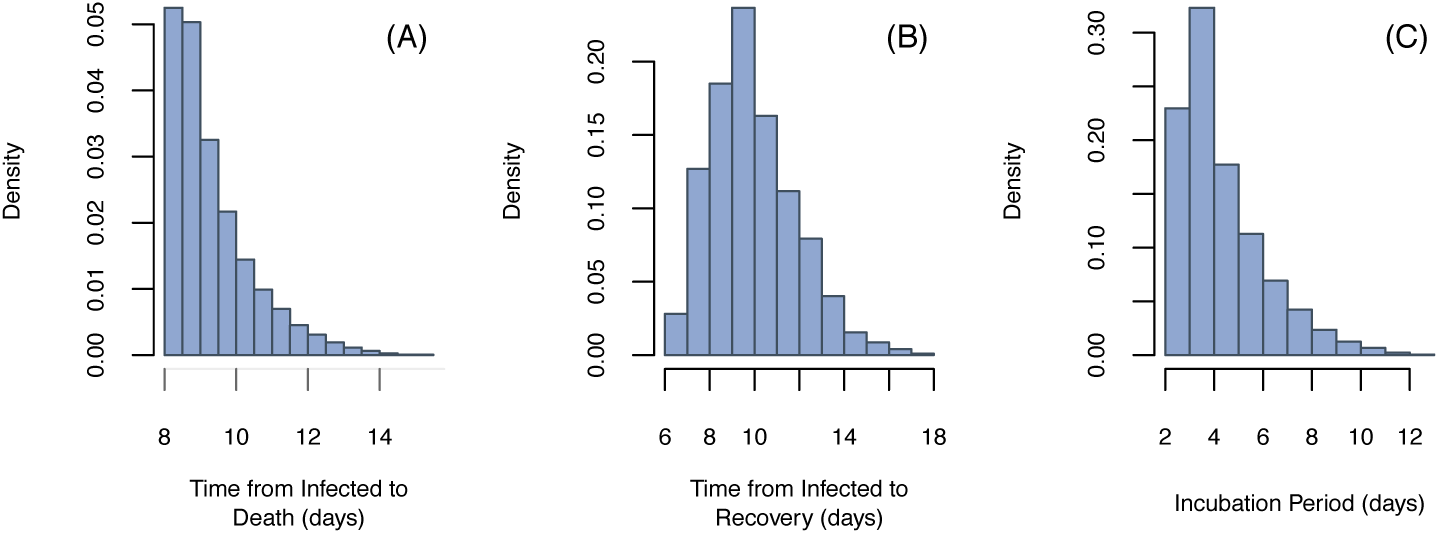
Extrapolations of the delay distributions post EBOV infection using within-host infection model. Simulations of the with-host infection model with varied infective dose and immune strengths. The median of the three distributions are 9, 9.7, and 3.8 for A, B, and C, respectively.

### The network model

Figure 4 shows an example of the generated networks and its required data. The network is returned as an adjacent matrix that is compatible to available network analyses algorithms, e.g., igraph [56]. Storing as a sparse matrix, a regular installation of R can generate reliably networks 10-20 thousand nodes with the generation time 6-10 minutes on a single thread Intel Core i7, 8GB RAM. In particular, giving a network of size *N* ∈ ℕ, each node is assigned an age such that the network’s age-distribution resemble that of a target population. Subsequently, nodes were assigned a number of contacts per day follow a defined contact distribution. Finally, the algorithm visits each node to generate the defined number of contacts, not at random but follow a defined contact matrix.

**Figure 4.**
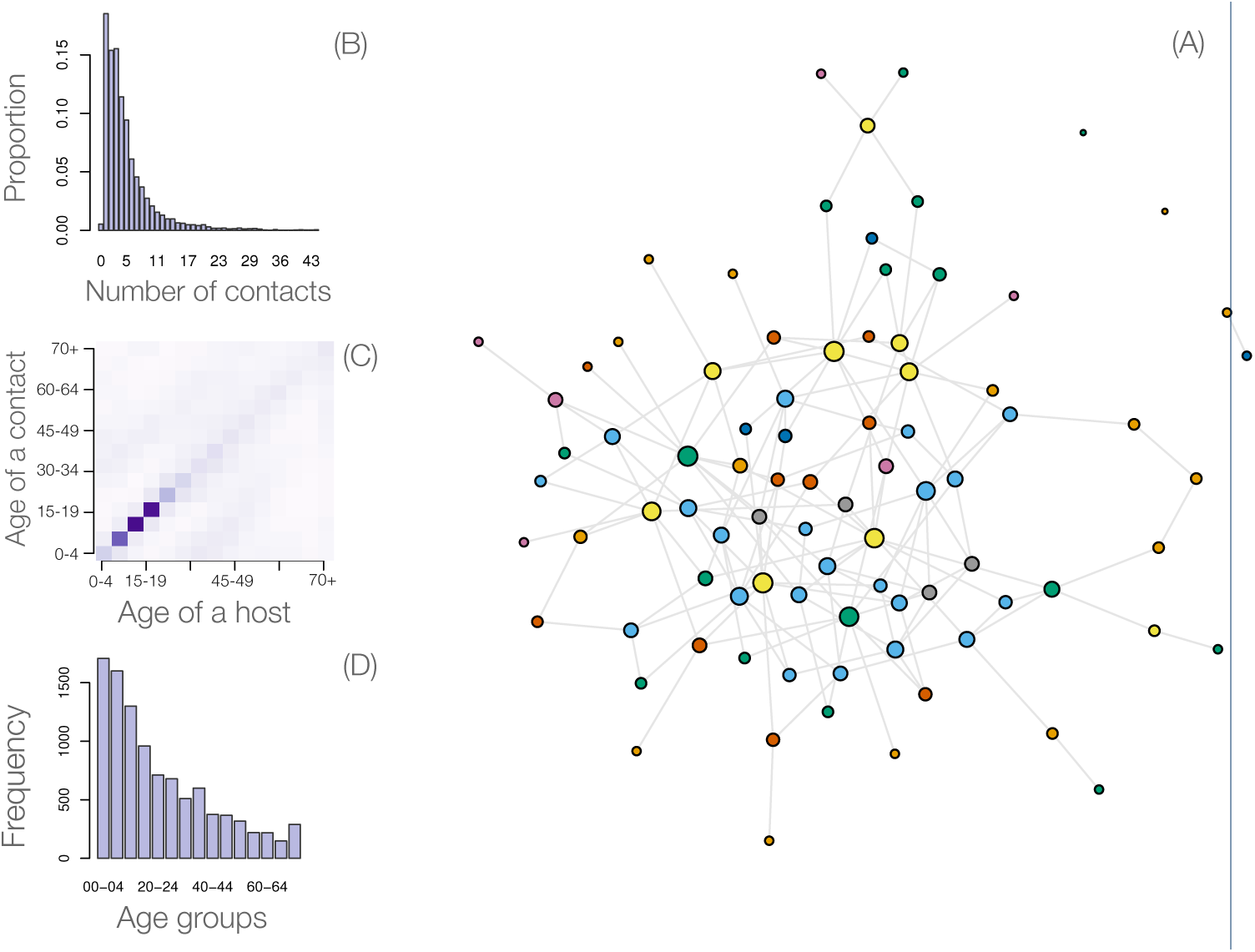
**A** Generated network of one hundred individuals that mimics distribution of physical contact, contact matrix, and population age structure. The node’s size reflects its number of contacts. Nodes in the same age-group have the same colour; **B** Distribution of number of physical contacts shows a majority of individuals have a few physical contact per day [31]; **C** A heat map of contact matrix shows higher contact frequencies in darker shades. The matrix reflects the assortative pattern of human contacts, that is people contact mainly with their peers, follow by their children or parents. The age-group with the highest contacts are teenager and young adults [31]; **D** Reconstructed age-structure of Sierra Leon population based on Statistics Sierra Leone and ICF International data [52].

### Calculating basic reproductive number (R0)

After each simulation, the uninfected nodes were removed from the initial network. Then the reproductive number is calculated as the average network degree, considering the network as directed and without loop (Fig. 5-Left). Simulation results showed that the overall estimate of the R0 was 1.43 (Fig. 5-Right). However, the estimates differed by age-groups with the highest of 4.7 for the group of 10-14 years of age. Generally, the age-groups with a higher contact rate had also a higher R0. Simulations of epidemics with varied intervention strategies showed that the Re can be reduced below one if the vaccination program with 85% coverage were deployed as far as five months before the introduction of the index case (time zero) or as late as one week after that (Supplemental Figure 1). This coverage threshold was tested as it is the highest vaccine coverage currently achieved worldwide for some diseases, e.g., Hepatitis B, measles, and polio [57]. Late initiations of similar interventions from one to five months after the time zero gradually shift the Re to the outbreak domain.

**Figure 5.**
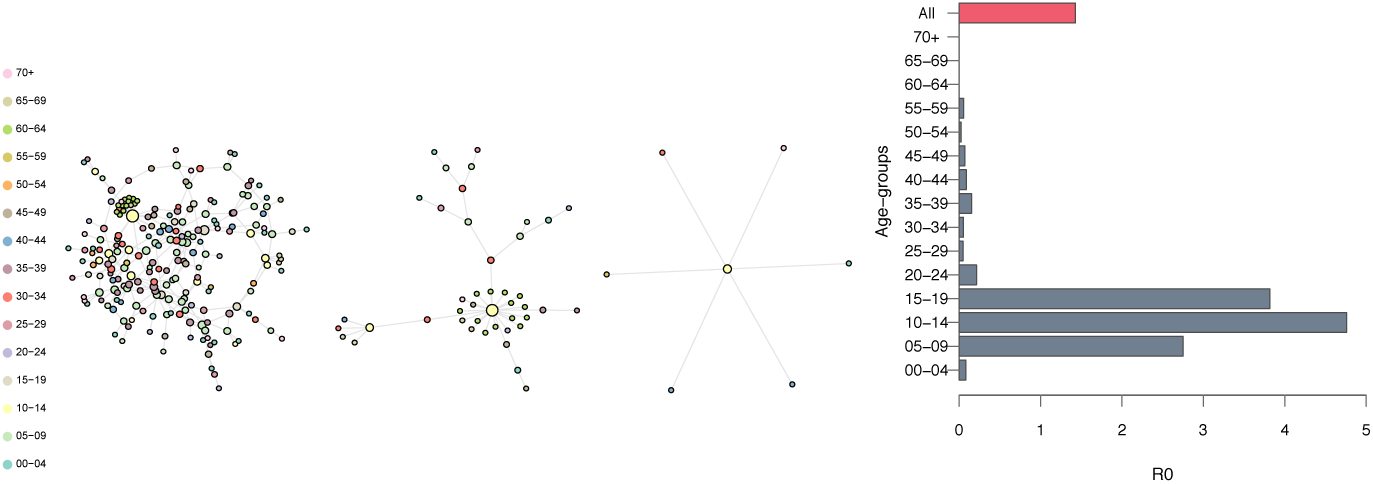
Left. Three examples infected networks. The three networks were randomly chosen from the simulated epidemics. Uninfected nodes were removed and the network are plotted. Based on these, the R0 was calculated based on the edges assuming a directed network, i.e., each edge counted in only one direction. **Right. Estimates of the basic reproductive number without any intervention, overall and by age-groups**. Simulations of a network of size ten thousand during a period of one year. One thousand simulations were run, each time with a random index case. At the end of each simulation, networks of infected nodes were extracted to compute the average number of secondary infections.

A lower vaccination coverage of 33% appeared not protective and posed a potential of outbreak regardless the time of vaccination program (Supplemental Figure 1). This coverage was tested as it is a theoretical protective threshold, i.e., 1-1/R0 [58]. Note that the tested time window of five months before the appearance of the index case was chosen based on the windows of opportunity for EBOV vaccination [38]. As of now, no data are available on the secondary antibody responses to EBOV; it was assumed that secondary responses are similar to the primary responses.

### Case-fatality rate

Simulations showed that the case-fatality rate in the absence of intervention is 90.93% (Supplemental Figure 2) which falls in the range of literature estimates of 0.4 to 0.91 [5]. Furthermore, simulation results showed that all the intervention strategies mentioned previously can reduce the case-fatality rate. These results highlight a benefit of vaccination programs even they are late, i.e., they can reduce the disease severity in newly infected subjects after the vaccination program. As such, relying solely on R0 as the indicator for evaluating intervention programs could have overlooked this life-saving aspect.

### Epidemic final size

Theoretical analyses of epidemic models showed when the R0 is larger than one, the final size of an epidemic will converge to a two points distribution: either the epidemic dies out with a small number of infected cases or the epidemic takes off and converges to a normal distribution [58]. Simulation results confirmed this epidemic behavior (Fig. 6). The results showed that without intervention, EBOV had approximately 50% to infect more than half the population. The introduction of vaccination programs at both the coverage thresholds previously mentioned and at any vaccination time points under assessments were able to scale down the epidemic size (Fig. 6). The two points epidemics size distribution gradually converged to a uni-modal distribution centring at a low infected fraction when the vaccination programs were deployed earlier. The high vaccine coverage strategy can effectively eliminate the possibilities of having a major outbreak infecting a large proportion of the population. This can be achieved when the vaccination programs were deployed any time from a week to five months before time zero.

**Figure 6.**
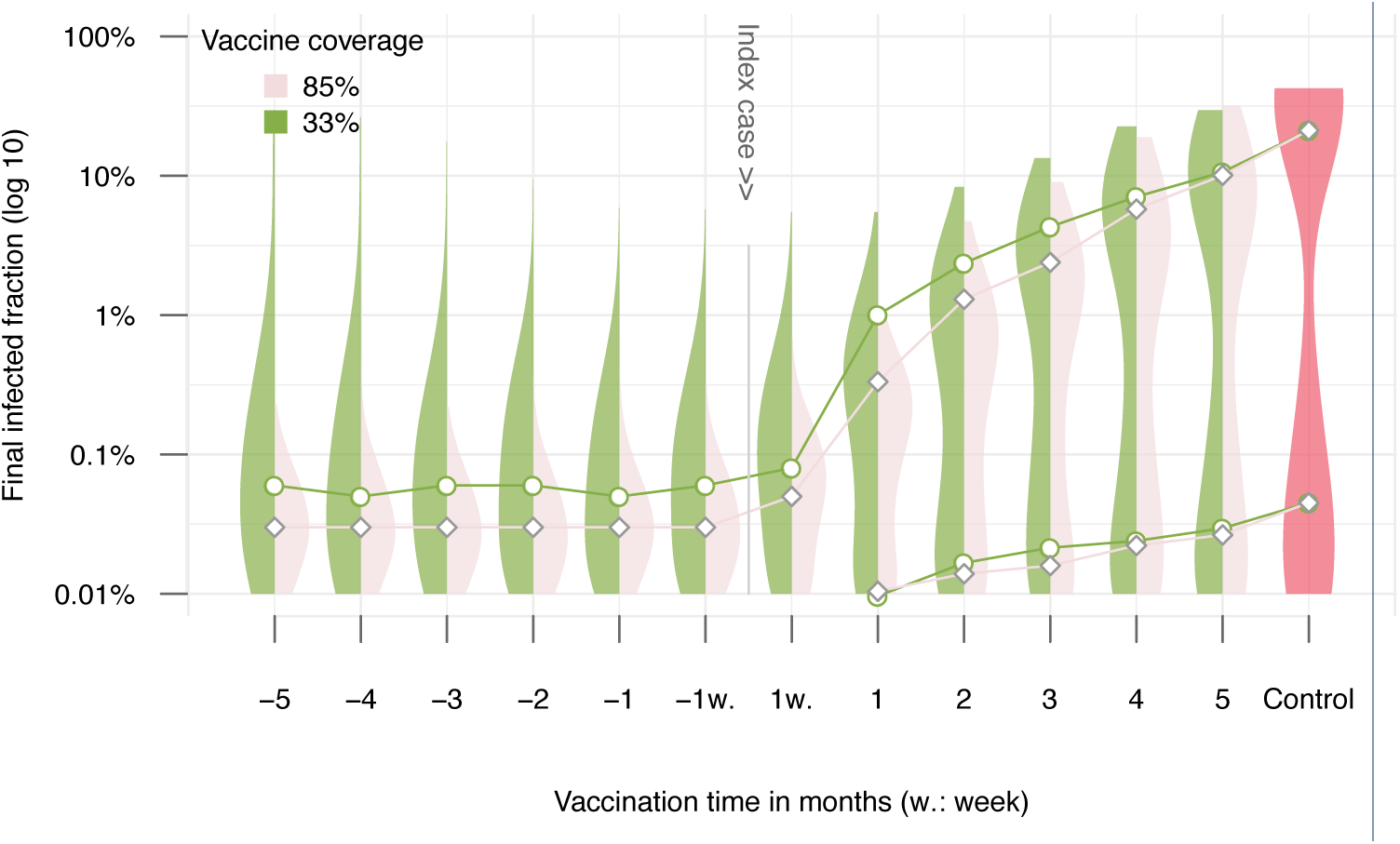
Distribution of the final infected fraction in different timing and coverage of vaccination strategies. A synthetic population of ten thousand individuals was generated. One thousand simulations were run to simulate the epidemic in the time course of one year. Each time, one individual was chosen randomly as the index case. Circles, diamonds, and connected lines are median. Filled areas are the corresponding non-parametric densities estimates [59]. Two median values are presented for multi-modal density estimates, determining by inflection points.

A random vaccination program covering 33% of the population one week before the epidemics can reduce the final size by more than 100 times compared to a no intervention scenario. However, the low coverage strategy still showed a small probability that epidemics can become major whereas the high coverage strategies did not. Vaccination programs deployed during the epidemics can also substantially reduce the epidemics size. The intervention conducted one month after time zero can also reduce the final size by more than ten times. These interventions not only able to reduce the final size, but they can also increase the epidemics extinction probability.

## Discussion

Epidemic modelling aims to obtain generalized solutions to questions such as whether or not a substantial population fraction is getting infected? how large would the outbreak spread? and how can the outbreak be mitigated with certain intervention approaches [58, 7]. Answering those questions requires the use of assumptive parameters as well as actual outbreak data [7, 58, 26, 15]. Our results showed that using information on within-host infection dynamics allows the identification of those key characteristics in the disease transmission.

Estimates of the incubation period suggest a contact tracing period of three weeks for Ebola epidemics, matching the current WHO’s recommendation of 21 days [60]. Estimates of the delay distributions agreed that EBOV infected subjects can be infectious from day 3 up to three weeks post infection [5]. Understanding of these delay distributions is critical in both clinical and epidemiological perspectives [61]. These distributions, however, are most often only partially observed in practice: it is difficult to know the exact time of exposure to the pathogen or to have complete outbreak data [6, 62]. As such, parameter estimation of these distributions have been relied on testing and comparing different distributional assumptions [62]. In this paper, mechanistically generated transmission characteristics using viral dynamics remarkably resemble literature estimates of Ebola. This approach is thus promising and practical given the accumulating experimental data on varieties of pathogens, notably, the one that as yet unknown in epidemic contexts.

To determine infection outcomes, the threshold AUC_7_ was chosen based on suggestions from empirical data in humans [49] and non-human primates [39, 40]. Simulations of the epidemics using this threshold revealed faithful estimates of the EBOV case-fatality rate (Supplemental Figure 2), supporting the use of the total viral load (AUC) as a criterion for determining infection outcomes. Although a more precise threshold criterion is desirable, it might not be feasible to obtain in practice considering inherent ethical reasons. Thus a similar criterion could be considered when adapting this approach to other infectious diseases, but ideally with dedicated experimental data.

Different classes of network models have been proposed, but they cannot reproduce properties observed in real world networks [63]. In addition, choices of theoretical network structure used for simulation can alter epidemic outcomes [28]. Thus, a network model at least obeying empirical data provides a more solid ground for epidemic simulations. Apart from mimicking the contact data properties, our network model can express age-related infection traits via the assigned age attributes. It was used here to express individual differences in the susceptibility to viral infection—an important element in a realistic disease transmission. Although contact data might not be available for a certain target area, the assortative patterns of human contacts and the highly skewed distribution of the number of contacts might hold true across regions [31, 32]. Thus, this paper presents a simple way to bring empirical data into epidemic modelling studies.

Our current network model currently can only simulate epidemics in a small population of size 10-20 thousand. This is because of the limit in R with the theoretical maximum square matrix size is approximately 45000 [55]. A more efficient storing of the network could extend the network size, such as a lazy evaluation used in igraph [56]. However, it could be more realistic to have several communities amount to a large population size instead of a large single network. This can be implemented by generating different communities across computers and allow them to communicate, speeding also the computation processes [64]. In this case, additional data are needed to model the communication among the communities, such as transportation and immigration.

Regarding EBOV epidemics, previous R0 estimates based on epidemic data were diverse, depending on model choices and assumptions [5]. Our estimate of R0 was 1.4 which is within the range of the previous estimates, ranging from 1.2 to 2.6, with some exceptional estimates up to 4.7 and 8.3 [5]. Notably, the estimates differed by age-groups with the highest of 4.7 for the group of 10-14-years of age. Although these estimates depend on Sierra Leon’s age-structure, the differences of R0 estimate stress the role of the age-structure and contact patterns in the estimation of R0, prompting that age-specific intervention strategies should be considered. The estimates by sub-groups single out the effort required to control the epidemic [9]. With current assumptions, targeting interventions to the group 5-20-years of age could be an effective strategy. Note that the differences of R0 by age-group could explain the wide variation of the previous estimates where different samples were employed [5].

The following assumptions were used in the paper given the lack of specific experimental data, but further efforts to produce data are needed to refine and to adapt to other settings: (i) Secondary antibody responses are the same as primary responses: This underestimates the effect of the vaccination strategies conducting before the epidemics. Experimental studies on secondary immune responses to EBOV infection are needed, especially those with a longer follow-up period. (ii) The transmission potential is directly related to viral load: This is although simple and reasonable, but different types of relationship, such as non-linear, might exist [13]. Dedicated animal experiments to define the exact relationship between the viral load the ability to transmit the virus are needed. (iii) The contact pattern is the same between European countries and Sierra Leone: Although the contact patterns seemed similar across countries [31], a more sociable population would increase the estimate of R0. (iv) Infection statuses have no influences on the network structure, except those were buried. This could overestimate R0 [65]. Taking people’s behaviour changes into epidemic modelling remains a grand challenge [65]. (v) Susceptibility to EBOV infection is similar to a general viral infection disease: Studies on susceptibility functions are lacking and require more attentions of the infection research community.

## Conclusion

Throughout this paper, we showed the possibilities to investigate practical and intriguing questions using a within-host viral dynamic model and an age-structured network model. The advantages of using explicitly within-host dynamics are the availability of experimental data, the possibility of conducting experiments to characterize transmission parameters, and the ability to provide high-resolution subject-specific responses to infection. The advantages of using an age-structured network model are its simple implementation, its representativeness for disease transmission, and the availability of the age-structured data. Therefore, immunological studies of infectious agents could be seamlessly integrated into studies of between hosts transmission, promoting evidence-based public health practices.

## List of abbreviations

EBOV: Ebola virus
NHPs: Nonhuman primates
VSV: Vesicular stomatitis virus
R0: Basic Reproduction Number
Re: Effective Reproduction Number
AUC: Area Under the Curve
WHO: World Health Organization

## Ethics approval and consent to participate

Not applicable.

## Consent to publish

Not applicable.

## Availability of data and materials

All data generated or analysed during this study are included in this published article and its supplementary information files.

## Author’s contributions

EAHV supervised the project. VKN designed the modelling and performed the simulations. VKN, EAHV and RM provided and analysed the data. VKN, EAHV and RM discussed and wrote the manuscript. All authors reviewed the manuscript.

## Acknowledgements

Not applicable.

## Additional Files

### Code and examples — Generating age-specific contact network

The R code for generating an example contact network can be previewed at http://doi.org/10.5281/zenodo.1037264

### Code and examples — Epidemic simulations

Example R code for epidemic simulations can be previewed at https://doi.org/10.5281/zenodo.1045404

#### Supplemental Figure 1

**Figure 7.**
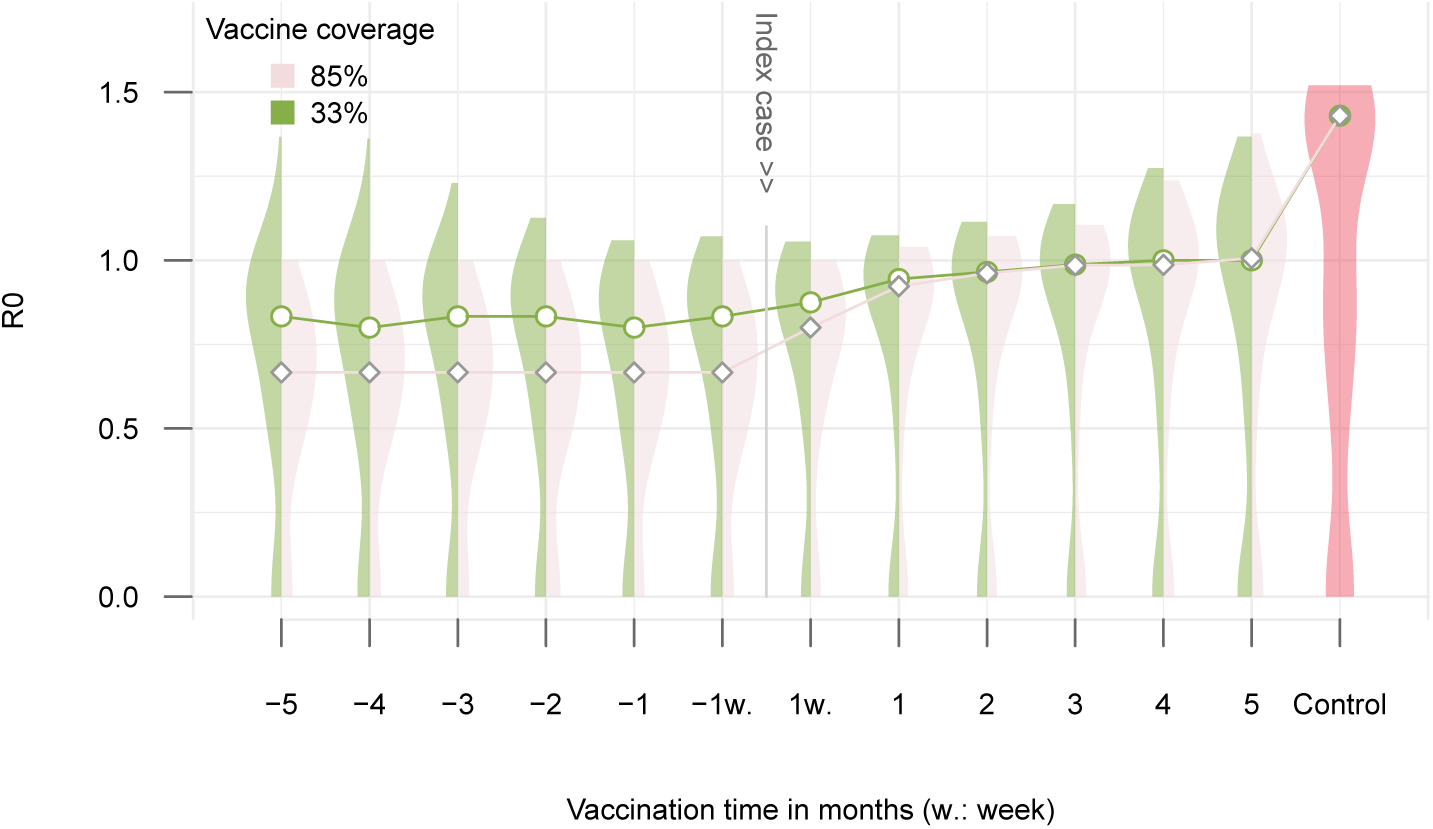
Estimates of the reproductive number in different vaccination schemes. Simulations of a network of size ten thousand during a period of one year. One thousand simulations were run, each time with a random index case. At the end of each simulation, network of infected nodes were extracted to compute the average number of secondary infection.

#### Supplemental Figure 2

**Figure 8.**
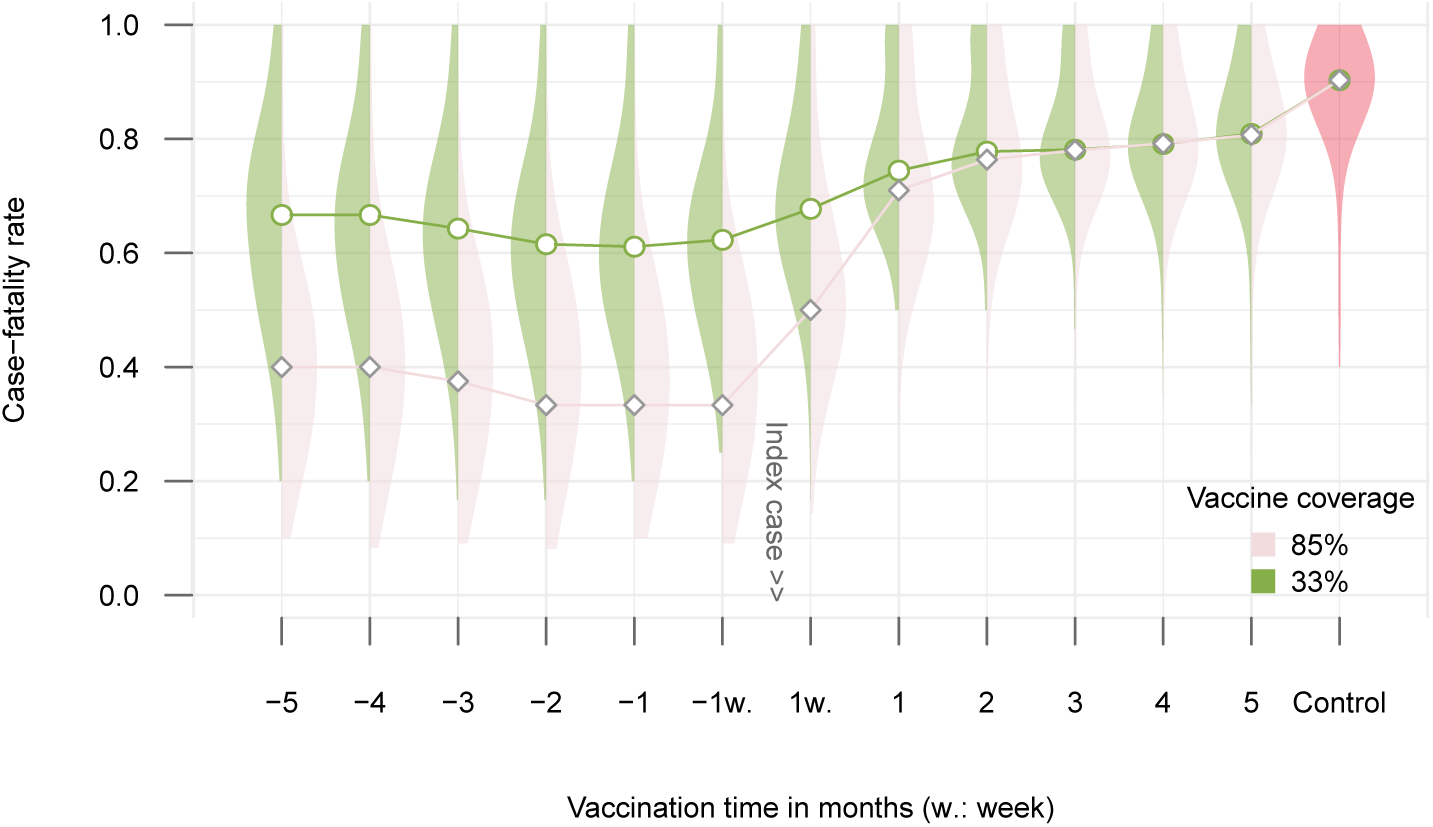
Case-fatality rate in different vaccination schemes. Simulations of a network of size ten thousand during a period of one year. One thousand simulations were run, each time with a randomly index case. At the end of each simulation, the network of infected nodes was extracted to compute the case-fatality rate.

